# Lamin A/C phosphorylation at serine 22 is a conserved heat shock response to regulate nuclear adaptation during stress

**DOI:** 10.1101/2022.01.19.476880

**Authors:** Laura Virtanen, Emilia Holm, Mona Halme, Gun West, Fanny Lindholm, Josef Gullmets, Juho Irjala, Tiina Heliö, Artur Padzik, Annika Meinander, John E. Eriksson, Pekka Taimen

## Abstract

The heat shock (HS) response is crucial for cell survival in harmful environments. Nuclear lamin A/C, encoded by *LMNA* gene, has been shown to contribute towards altered gene expression during heat shock, but the underlying mechanisms are poorly understood. Here we show that reversible lamin A/C phosphorylation at Ser22 upon HS is an evolutionary conserved stress response that is triggered in concert with HSF1 activation in human and mouse cells and can also be observed in *D. melanogaster in vivo*. Consequently, the phosphorylation increase facilitated nucleoplasmic localization of lamin A/C and nuclear rounding in response to HS. The importance of lamin phosphorylation equilibria in HS was confirmed by lamin A/C knock-out (KO) cells that showed deformed nuclei after HS and were rescued by ectopic expression of wild-type, but not by a phosphomimetic (S22D) lamin A mutant. Furthermore, HS triggered release of lamina-associated protein 2α (Lap2α) from its association with lamin A/C and concurrently its downregulation, a response that was perturbed in lamin A/C KO cells and in *LMNA* mutant patient fibroblasts. The abrogated Lap2α response resulted in impaired cell cycle arrest under HS and compromised survival at the recovery. Taken together, our results suggest that the altered phosphorylation stoichiometry of lamin A/C provides an evolutionary conserved mechanism to regulate lamin structure and serve nuclear adaptation and cell survival during HS.

## INTRODUCTION

The heat shock (HS) response is an evolutionally conserved mechanism and vital for cell survival in harmful environments or pathophysiological stresses such as high temperature, fever, or inflammation (Åkerfelt et al., 2010). Exposure to stressful conditions leads to synthesis of heat shock proteins (HSPs) that are in a key role to protect cellular homeostasis (Lindquist, 1986). The expression of HSP genes during HS is mainly controlled by heat shock factor 1 (HSF1) (Wu, 1995). Upon stress, inactive HSF1 monomers are converted to an active trimer that is hyperphosphorylated and translocated into the nucleus. Active HSF1 binds to heat shock elements (HSE) at the promoter regions of HSP genes and induces transcription (Åkerfelt et al., 2010). Transcriptional response to HS results in the activation of hundreds and repression of thousands genes. HSF1 is critical for the induction of HSPs and over 200 other genes; however, the induction and repression of transcription during HS are comprehensive and the majority of these changes are HSF1-independent (Mahat et al., 2016). HSF1 DNA binding activity is detected within minutes of HS and is maintained at a high level for an hour. Thereafter, HSF1 DNA binding activity decreases to control levels, even if the cells still remain exposed to the HS (Kline & Morimoto, 1997).

Accumulating data show that the nuclear lamina is dynamically remodeled in response to environmental cues. The ratio of A-type lamins (primarily lamin A and C) and B-type lamins (lamin B1 and B2) correlates with tissue stiffness, and the elasticity and organization of the lamina is further regulated by lamin phosphorylation/dephosphorylation status (Kochin et al. 2014; Buxboim et al., 2014; Swift et al., 2013). There are a number of studies suggesting that that altered dynamics of lamins contribute to and are beneficial for cellular adaptation under HS. Early studies reported heat-induced stabilization of nucleoskeleton and dephosphorylation of lamin A/C during HS (Krachmarov & Traub, 1993). Several studies have demonstrated that lamin B1 also is upregulated upon HS (Dynlacht et al., 1999; Pradhan et al., 2020; Zhu et al., 1999) and downregulated during recovery (Haddad & Paulin-Levasseur, 2008). In *Drosophila* Schneider 2 cells, lamin Dm2 is dephosphorylated to lamin Dm1 during HS (Smith et al., 1987). Furthermore, hypersensitivity to HS has been reported in dermal fibroblasts obtained from Hutchinson-Gilford progeria (HGPS) and familial partial lipodystrophy (FPLD) patients carrying G608G and R482Q/W mutations in the lamin A/C gene (*LMNA*), respectively (Paradisi et al., 2005; Vigouroux et al., 2001). Interestingly, a recent study demonstrated that HS upregulates lamin A/C, and that lamin A/C are required for heat-shock-mediated transcriptional induction of the Hsp70 genes (Pradhan et al., 2020).

As lamin A/C has been shown to contribute to the conserved HS response, we wanted to explore whether this response could be coupled to lamin A/C phosphorylation, which we have shown to be a primary determinant of lamin organization and assembly in interphase cells (Kochin et al., 2014). We show that lamin A/C is phosphorylated at serine 22 upon prolonged HS in both transformed and primary human cell lines, as well as in mouse fibroblasts, and *Drosophila melanogaster in vivo*. Lamin A/C phosphorylation leads to increased nucleoplasmic localization of lamin A/C and rounding of the nucleus in human fibroblasts. Furthermore, data from lamin A/C knock-out (KO) and mutant cells demonstrated that a balanced lamin structure is needed to avoid nuclear deformation under HS. In addition, our results reveal that lamin A/C is required during prolonged HS to regulate the functions of the nuclear lamin A/C binding partner, lamina-associated polypeptide 2 alpha (Lap2α), which we show to determine cell cycle progression and cell survival during HS.

## RESULTS

### Lamin A/C is phosphorylated at serine 22 during HS

In an attempt to gain a better understanding of the function of nuclear lamina under heat-induced stress, we examined the expression levels, localization, and post-translational modifications of nuclear lamin A/C during HS. To this end, HeLa cells and primary human and mouse fibroblasts were exposed to HS for different periods of time (Fig. 1A). We used HSF1 gel shift as an established marker for an indicuded HS response (Åkerfelt et al., 2010). Intriquingly, in concert with induced HS response, we observed gradually increased phosphorylation of lamin A/C at serine 22 residue (Ser22) upon 1- to 4-hour HS at 42°C in all the cell lines (as normalized to lamin A/C and GAPDH) while phosphorylation of serine 392 remained relatively stable throughout the experiments (Fig. 1A, Fig. S1). Whole *D. melanogaster* fruit flies heat-shocked for 30 minutes at 37°C showed similarly increased phosphorylation at Ser37, which is homologous to Ser22 in mammals (Fig. 1A). During the recovery phase, pSer22 levels were reduced along with the attenuation of the HS response as indicated with the lost gel shift of HSF1 (Fig. 1A, Fig. S1).

**Figure 1.**
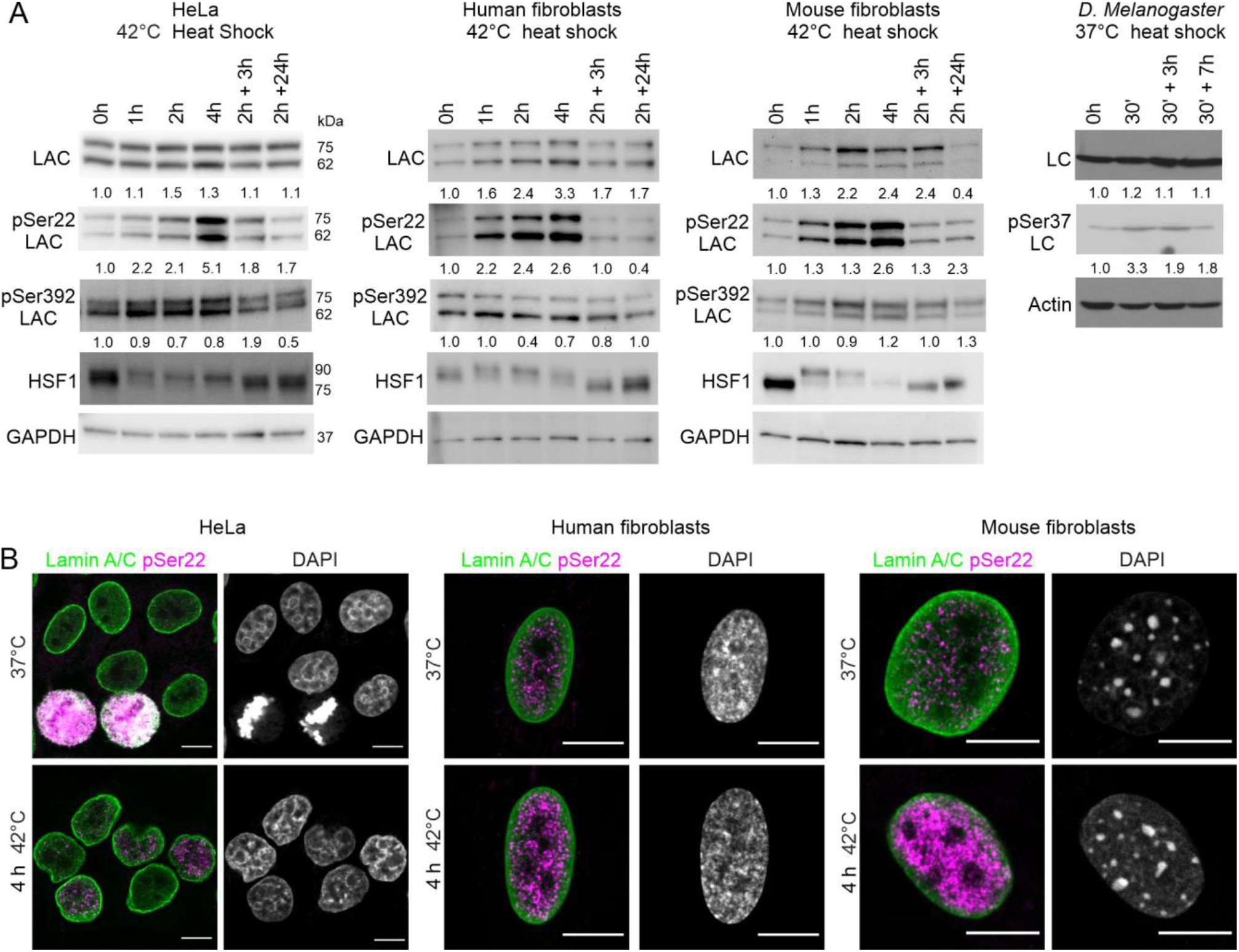
Lamin A/C is phosphorylated at serine 22 upon HS. **A)** Western blot analysis of lamin A/C (LAC), phospho-serine 22 lamin A/C (pSer22 LAC), phospho-serine 392 lamin A/C (pSer392 LAC), and heat shock factor 1 (HSF1) after 1- to 4-hour heat shock (HS) at 42°C and 3- and 24-h recovery at 37°C in Hela cells, primary human and mouse fibroblasts. Whole *D.Melanogaster* flies were heat-shocked for 30 min at 37°C and left to recover for 3 and 7 h at 22°C. The average numerical values of signal intensities relative to loading control (GAPDH, actin) from individual experiments are shown below each blot. pSer22 and pSer392 lamin A/C levels were normalized to GAPDH and lamin A/C. (Hela: N=6, human fibroblasts: N=3, mouse fibroblasts: N=3, *D.Melanogaster:* N=4). **B)** Hela cells, human and mouse fibroblasts were cultured either in normal culture conditions or exposed to 4-h HS at 42°C, fixed and stained for lamin A/C (green), pSer22 lamin A/C (magenta), and DAPI (grey). Scale bar 10 μm.

Additionally, we carried out a mass spectrometry (LC-MS/MS) analysis to identify any additional differently phosphorylated residues in lamin A/C. Apart from Ser22, there were five residues, Thr3, Ser5, S268, Ser398, and Thr590, that were detected in HS samples but not in control samples (Table S1). These sites correspond well to the primary interphase sites that we have previously shown to be primarily involved in regulating lamin A/C assembly and dynamics (Kochin et al., 2014). To check how the HS-mediated Ser22 phosphorylation relates to the previously identified mitotically-induced Ser22 phosphorylation, HeLa cells were arrested at G1/S phase with aphidicolin prior to HS. Lamin A/C was equally phosphorylated in synchronized cells upon HS confirming that phosphorylation of lamin A/C at Ser22 takes place in heat-shocked interphase cells rather than in mitotic cells (Fig. S1).In comparison, the nocodazole-arrested mitotic control cells had significantly more pSer22 lamin A/C compared to the heat-shocked cells at G1/S phase (Fig. S1). Confocal microscopy analysis with pSer22 lamin A/C antibody also showed increased nucleoplasmic labeling in non-mitotic human and mouse cells upon HS (Fig. 1B). This analysis also showed that the phosphorylation level is significantly higher in mitotic cells where all the lamin is in a disassembled nucleoplasmic state. While the HS-induced phosphorylation is at a lower level, the shift from the assembled form at the nuclear lamina to nucleoplasmic lamin is also more modest. These results indicate that lamin A/C phosphorylation at Ser22 is an evolutionary conserved HS response mechanism the onset and cessation of which occur in concert with HSF1 activation and attenuation.

### Phosphorylation of lamin A/C facilitates rounding of the nucleus in response to HS

To study further how HS and lamin phosphorylation affects nuclear and lamina structure, primary human dermal fibroblasts from a healthy individual and from a dilated cardiomyopathy (DCM) patient carrying the p.S143P mutation in *LMNA* were analyzed upon mild 42°C (Fig. 2A) and severe 44°C HS (Fig. 2B). The baseline pSer22 status was slightly higher in the patient cells compared to controls (Fig. 2A, B; p < 0.05). While the HS-induced increase in lamin A/C phosphorylation followed the same kinetics in both cell lines, the pSer22 levels remained consistently higher in the patient cells than in the control cells (Fig. 2A), an effect that was especially conspicuous upon severe 44°C HS (Fig. 2B). Noteworthy is that the lamin A/C phosphorylation increased several fold higher in cells exposed to severe HS as compared to cells subjected to mild HS (Fig. 2A, B). Following HS at 42°C (4h) and 44°C (1h), there was a slight but statistically significant nucleoplasmic shift of lamin A/C in the control cells (Fig. 2C, D; p < 0.01). In comparison, when patient fibroblasts were examined, lamin A/C was found to be more nucleoplasmic already under normal culture conditions (Fig. 2C) corresponding to our previous observations (West et al., 2016), and the HS-induced increase in nucleoplasmic lamin A/C labeling was detected only at 44°C (Fig. 2C, D; p < 0.05). Correspondingly, lamin A/C phosphorylation intensity correlated with increased nucleoplasmic localization when analyzed from individual cells at 37°C and upon 1- to 2-h HS at 44°C (Fig. 2E; Pearson correlation coefficient R=0.38 for control and R=0.35 for patient cells, p < 0.01).

**Figure 2.**
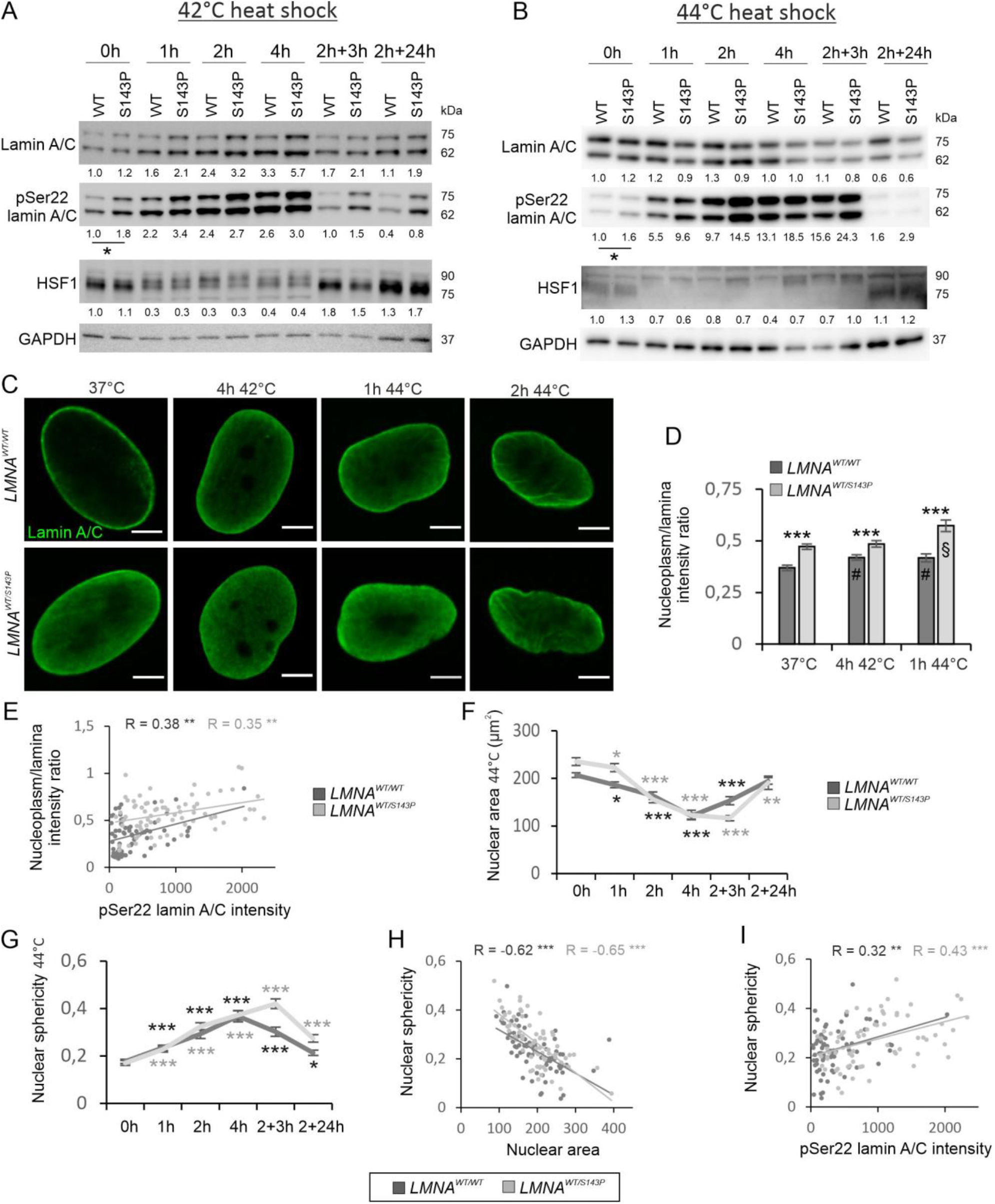
Phosphorylation of lamin A/C facilitates rounding of the nucleus in response to HS. **A-B)** Western blot analysis of control and patient fibroblasts carrying the p.S143P mutation in *LMNA* as detected with lamin A/C, pSer22 lamin A/C and HSF1 antibodies upon 1- to 4-h HS at 42°C or 44°C and at the recovery. The average numerical values of signal intensities relative to the loading control (GAPDH) are shown below each blot (N=3-5 replicates). pSer22 lamin A/C was normalized to GAPDH and lamin A/C. **C)** Confocal microscopy images of *LMNA^WT/WT^* and *LMNA^WT/S143P^* fibroblasts stained for lamin A/C at normal culture conditions, after 4-h HS at 42°C and after 1-h and 2-h HS at 44°C. Scale bar 5 μm. **D)** Lamin A/C fluorescence intensities at the lamina region and in the nucleoplasm were determined from the mid-plane confocal sections of randomly selected cells and the average ratios of the signals (nucleoplasm/lamina) were plotted (N=30-50). **E**) Scatter blot of lamin A/C nucleoplasm/lamina intensity ratio versus pSer22 lamin A/C intensity determined from randomly selected cells at 37°C and after 1- and 2-h HS at 44°C (N=70). **F-G)** Nuclear area (μm^2^) and nuclear sphericity of *LMNA^WT/WT^* and *LMNA^WT/S143P^* fibroblasts was determined from confocal sections of randomly selected cells at 37°C, after 1- to 4-h HS at 44°C, and after 3- and 24-h recovery at 37°C (N=50-90). **H)** Scatter blot of nuclear sphericity versus nuclear area upon 0 - 2 h HS at 44°C (N=70). **I)** Scatter blot of nuclear sphericity versus pSer22 lamin A/C intensity upon 0 - 2 h HS at 44°C (N=70). Bars express mean ± s.e.m, Pearson correlation coefficients (R) are indicated, *p < 0.05, **p < 0.01, ***p < 0.001, # p < 0.01 (*LMNA^WT/WT^* control vs. *LMNA^WT/WT^* HS), § p < 0.05 (*LMNA^WT/S143P^* control vs. *LMNA^WT/S143P^* HS).

Next, we investigated whether lamin A/C phosphorylation status correlates with HS-induced morphological nuclear changes in individual cells. Determination of nuclear area and sphericity from confocal sections of lamin A/C stained cells releaved no significant changes under mild 42°C HS (Fig. S2). However, the nuclear area decreased significantly in both cell lines under 44°C HS (Fig. 2F) and inversely correlated with nuclear rounding (sphericity) (Fig. 2G-H; Pearson correlation coefficient R=−0.62 for control and R=−0.65 for patients cells, p < 0.001). Equally, lamin A/C phosphorylation intensity and nuclear sphericity correlated with each other in individual heat-shocked cells (Fig. 2I; Pearson correlation coefficient R=0.32 for control and R=0.43 for patients cells, p < 0.001). These results confirm that phosphorylation of lamin A/C facilitates nucleoplasmic localization of lamin A/C, which then results in rounding of the nucleus in response to HS.

### Lamin A/C phosphorylation at Ser22 is mediated by several kinases under HS

To identify potential kinase(s) responsible for lamin A/C phosphorylation during HS we tested whether aforementioned HS response could be prevented in HeLa cells when treated by different kinase inhibitors for 24 h prior to exposure to HS (Fig. 3A). In the presence of specific PKC (10 μM Go6976) and MAPK (10 μM U0126) inhibitors, no increase in lamin A/C phosphorylation was detected upon HS suggesting that both kinases may be involved in HS-induced lamin phosphorylation. Additionally, MAPK inhibited cells showed significantly less phosphorylation under HS compared to untreated HS cells (p < 0.05). Similarly, the generic kinase inhibitor staurosporine (STA 200nM; inhibitor of multiple kinases, e.g. PKC, MAPK) reduced lamin phosphorylation when compared to unheated cells and heat-shocked control cells (p < 0.05 and p < 0.01 respectively). AKT inhibition (10 μM API-2) decreased the phosphorylation under normal culture conditions, but lamin A/C phosphorylation was still slightly increased under HS. CDK inhibitors (100 nM flavopiridol and 1 μM roscovitine) had no effect on lamin A/C phosphorylation under HS.

**Figure 3.**
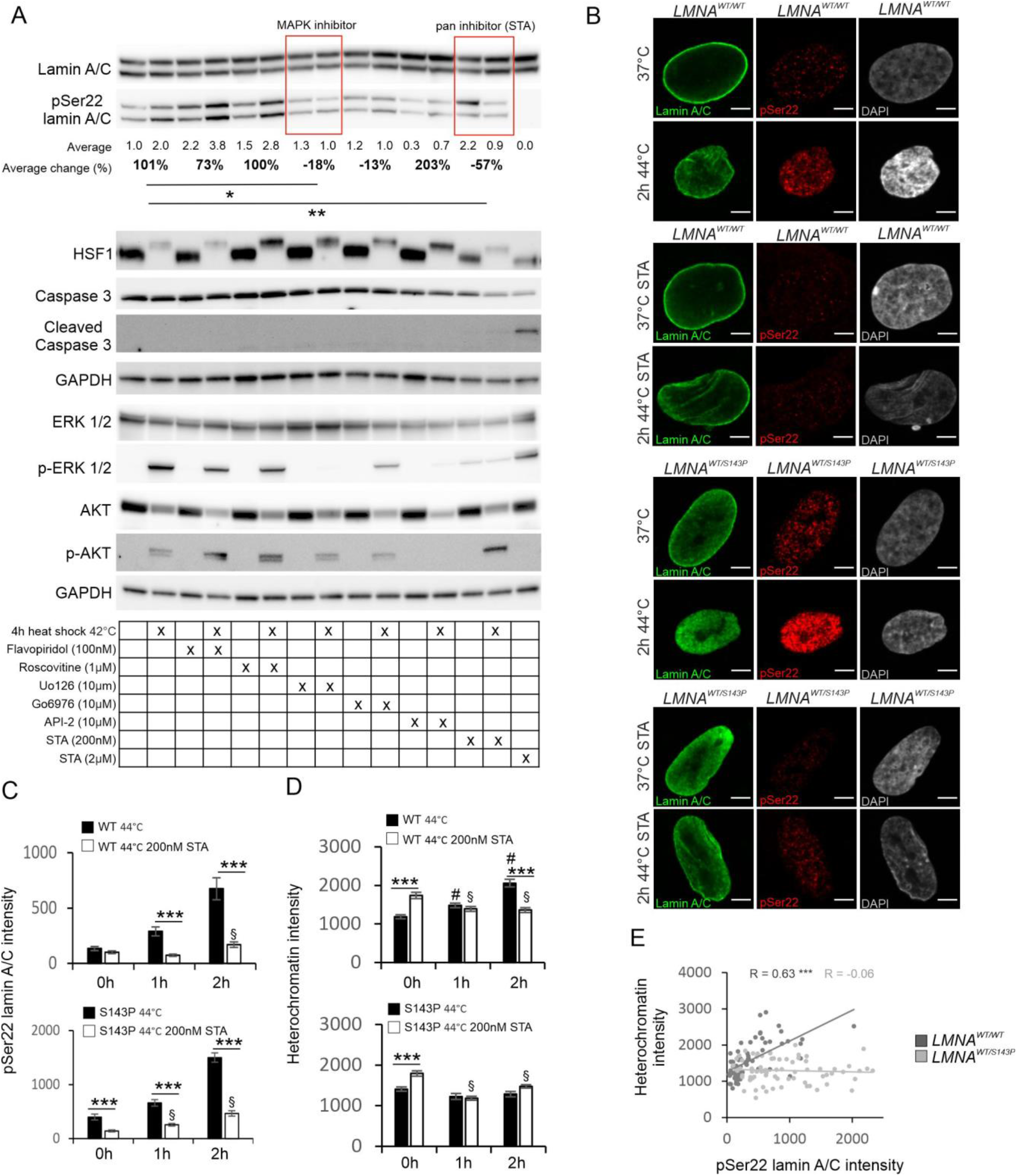
The effect of kinase inhibition on lamin A/C phosphorylation under HS. **A)** Immunoblots showing lamin A/C, pSer22 lamin A/C, HSF1, ERK1/2, pERK1/2, AKT, pAKT, caspase-3, and cleaved caspase-3 protein levels in control and heat-shocked HeLa cells treated with different kinase inhibitors. GAPDH was used as a loading control and HeLa cells treated with 2 μM staurosporine as a positive control for apoptotic cell death. The average numerical values of signal intensities relative to the loading control (GAPDH and lamin A/C) and percentage of intensity change between control and heat shocked cells are shown below the blot (N=5). **B)** Confocal images of *LMNA^WT/WT^* and *LMNA^WT/S143P^* fibroblasts stained for lamin A/C (green), pSer22 lamin A/C (red) and DAPI (grey). The cells were cultured either at 37°C or heat-shocked for 1 hour at 44°C in the absence or presence of 200 nm kinase inhibitor STA. **C-D)** Intensity values of pSer22 lamin A/C and heterochromatin (DAPI) were analyzed from confocal images of *LMNA^WT/WT^* (top two graphs) and *LMNA^WT/S143P^* (lower two graphs) cells. The cells were heat shocked for 1-2 hours at 44°C in the absence or presence of 200 nm STA (N=30). **E)** Scatter blot of heterochromatin (DAPI) intensity versus pSer22 lamin A/C intensity determined from randomly selected cells at 37°C and after 1- and 2-h HS at 44°C (N=70). Bars express mean ± s.e.m, Pearson correlation coefficients (R) are indicated, *p < 0.05, **p < 0.01, ***p < 0.001, ^#^p < 0.001 (control vs. HS), ^§^p < 0.01 (STA control vs. STA HS).

The effect of 200 nM STA on nuclear structure of control and patient fibroblasts was further analyzed after 1-h HS at 44°C. STA treatment reduced the pSer22 lamin A/C labeling intensity significantly in both cell lines although a minor phosphorylation increase was still detected under HS, especially in the patient cells (Fig. 3B, C; p < 0.001). Interestingly, STA treatment appeared to inhibit the morphological changes previously detected in heat-shocked cells. More specifically, the area and sphericity of heat-shocked healthy control cells remained unchanged in the presence of STA while heat-shocked patient cells still showed decreased nuclear area and rounding at 2-h timepoint (Fig. S3; p < 0.001). Similarly, an increase in heterochromatin intensity (as detected by DAPI) was inhibited by STA in healthy control cells after HS (Fig. 3D; p < 0.05) and correlated with lamin A/C phosphorylation status (Fig. 3E; p < 0.001). Surprisingly, similar increase or correlation was not detected in the patient cells (Fig. 3D, E). To conclude, these results suggest that several kinases are responsible, either directly of indirectly, for HS-induced lamin A/C phosphorylation. Since HS-induced increase in heterochromatin intensity is impaired in lamin mutant patient cells, these results also imply that defects in lamina structure may alter chromatin organization and global gene repression under HS.

### Lamin A/C knock-out cells exhibit deformed nuclear shape upon HS

To further study the role of lamin A/C under HS, we created stable lamin A/C knock-out (KO) HeLa cell lines using CRISPR/Cas9 technology. Cells lentivirally transduced with two separate sgRNAs showed no detectable amounts of lamin A/C after single-cell sorting, while transduction with non-targeting (NT) sgRNA had no effect on lamin expression (Fig. 4A-C). There was no significant difference in the nuclear area between the KO, NT and non-transduced parental cells under normal culture conditions or after 4-h HS at 42°C (data not shown). However, lamin A/C KO cells showed more convoluted nuclear shape already under normal culture conditions (Fig. 4D, p < 0.001) and these nuclear deformations became more evident upon HS whereas control cells retained their normal round shape (Fig. 4D, E; p < 0.001). The ectopic expression of wild-type (WT), phosphomimetic (Ser to Asp, S22D) and phosphorylation-deficient (Ser to Ala, S22A) mutant forms of GFP-labelled lamin A in LAC KO1 cells rescued symmetrical nuclear shape under normal culture conditions (Fig 4F-G; p < 0.001). However, only WT-LA and S22A-LA were able to protect nuclear morphology under HS (Fig. 4G, p < 0.001). As S22D-LA is mostly nucleoplasmic (Fig. 4F), this observation indicates that intact lamina structure is needed to support nuclear shape under HS.

**Figure 4.**
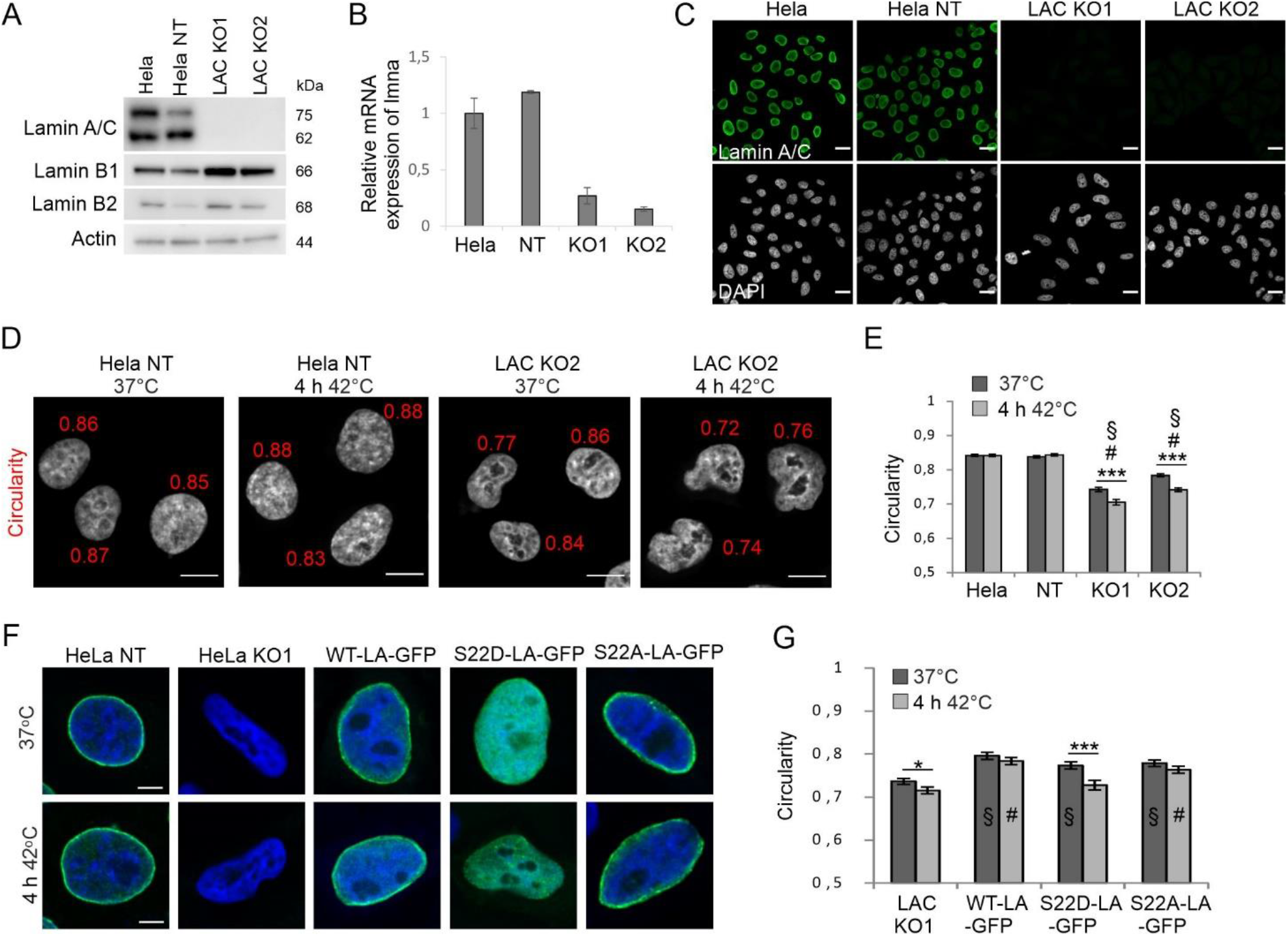
Knockout of the *LMNA* gene increases nuclear deformability under normal and HS conditions. **A)** Western blot analysis showing the expression of lamin A/C, lamin B1, and lamin B2 in parental Hela cells and cells treated with non-targeting (NT) gRNA or two different lamin A/C targeting gRNAs (LAC KO1 and LAC KO2). **B)** Expression of *LMNA* mRNA in parental Hela, NT Hela, LAC KO1 and LAC KO2 cells. **C)** Immunofluorescence images of parental Hela, NT Hela, LAC KO1 and LAC KO2 cells stained with lamin A/C (green) and DAPI (gray). Scale bar 20 μm. **D)** Representative images from nuclear circularity analysis of NT and KO2 cells at 37°C and after 4-h HS at 42°C. **E)** Nuclear circularity of parental Hela, NT Hela, LAC KO1 and LAC KO2 cells under normal and HS conditions (N=200, from 4 different experiments). **F)** Representative images of HeLa NT cells, untreated HeLa KO1 cells and HeLa KO1 cells transfected with WT-LA-GFP, S22D-LA-GFP or S22A-LA-GFP at normal culture conditions and after 4-h HS at 42°C. HeLa NT and HeLa KO1 were stained for lamin A/C (green). All cells were stained with DAPI (blue). **G)** Nuclear circularity of LAC KO1, WT-LA-GFP, S22D-LA-GFP and S22A-LA-GFP cells under control condition and after 4-h heat shock at 42°C (N=100). Data is expressed as mean ± s.e.m, *p < 0.05, ***p < 0.001, ^#^p < 0.001 (compared to HeLa and NT HeLa), ^§^p < 0.001 (compared to LAC KO1 control) ^¤^p < 0.001 (compared to LAC KO1 HS).

### HSF1 silencing affects lamin A but not lamin C protein level

Since HSF1 is known to influence protein expression during cell stress, we further studied whether HSF1 plays any role in the regulation of lamin A/C expression. HeLa cells stably silenced for HSF1 (shHSF1) were exposed to HS for pre-determined periods of time (Fig. 5A). Interestingly, we noticed an overall reduced expression of lamin A but not lamin C in shHSF1 cells suggesting that HSF1 may be involved in regulating lamin A protein level (Fig. 5A; N=5; p < 0.05; p < 0.01). Additionally, phosphorylation of lamin A/C at Ser22 appeared delayed in the shHSF1 Hela cells upon HS (Fig. 5A) but more detailed analysis showed that there was ~40% less pSer22 lamin A/C in shHSF1 cells already under normal conditions (as normalized to lamin A/C and HSC70) and the gradual increase in lamin A/C pSer22 level thereafter was similar to Hela WT (Fig. 5A). Immunofluorescence analysis showed no clear difference in lamin A/C staining in shHSF1 Hela cells compared to Hela WT (Fig. 5B). We also analyzed whether lamin A/C KO affects HSF1 protein expression but found no significant differences in HSF1 expression level or localization in the KO cells (Fig. S4). In summary, these results suggest that HS-induced phosphorylation of lamin A/C Ser22 is not HSF1-dependent although the activation and attenuation of HSF1 is closely linked to pSer-22 onset and cessation. However, lamin A downregulation in shHSF1 cells may indicate an indirect crosstalk between lamin A and HSF1.

**Figure 5.**
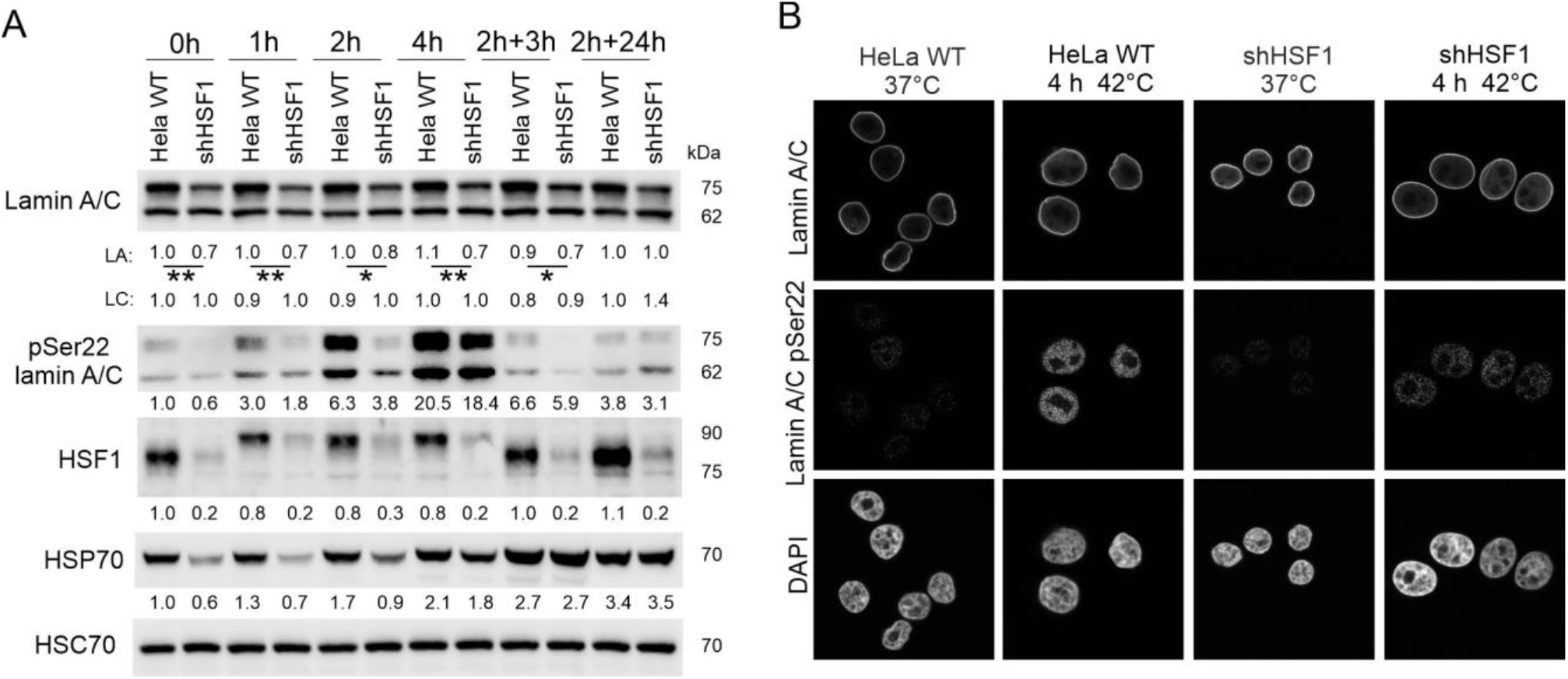
Lamin A/C phosphorylation at Ser22 is independent of HSF1. **A)** Western blot analysis of lamin A/C, pSer22 lamin A/C, heat shock factor 1 (HSF1), and heat shock protein 70 (HSP70) upon HS and after 3- and 24-h recovery. The average numerical values of signal intensities relative to loading control (HSC70) are shown below each blot. pSer22 lamin A/C levels were normalized to HSC70 and lamin A/C (N=5). **B)** Confocal microscopy images of parental HeLa cells and HSF1-silenced cells stained for lamin A/C, pSer22 lamin A/C and DAPI under normal culture conditions and after 4-h HS. Scale bars: 10 μm. * p < 0.05, **p < 0.01.

### Lap2α is degraded under HS in a lamin A/C dependent manner

We next asked whether lamin A/C phosphorylation affects its interaction with known binding partners. Since the phosphorylation mostly appears in the nucleoplasmic pool of lamin A/C rather than at the lamina region (Fig.1B), we decided to analyze the crosstalk between lamin A/C and its binding partner Lap2α. Based on proximity ligation assay (PLA), Lap2α and lamin A/C close proximity signals were reduced in HeLa cells after 4-hour HS at 42°C when compared to control cells (Fig. 6A, B; p < 0.001). Since decreased number of PLA signals may be due to reduced proximity or downregulation of either protein, we further tested the abundance of lamin A/C and Lap2α during HS. While lamin A/C protein levels remained relatively stable, there was a clear downregulation of Lap2α upon HS in parental Hela and NT Hela cells but interestingly not in LA/C KO1 or KO2 cells (Fig. 6C, S5). All the heat-shocked cell lines showed distinct Lap2α aggregates, which mostly appeared in the nuclear periphery (Fig. 6D). However, the Lap2α aggregates in the KO cells were approximately 300 nm closer to the center of the nucleus compared to control cells suggesting that loss of lamin A/C affected their localization (Fig. S5). The HS-induced Lap2α degradation was restored with ectopic expression of either WT-LA, S22D-LA, or S22A-LA into LA/C KO1 cells (Fig. 6E, S6). Additionally, the localization of Lap2α aggregates towards the nuclear periphery was rescued with all the plasmids (Fig. S6).

**Figure 6.**
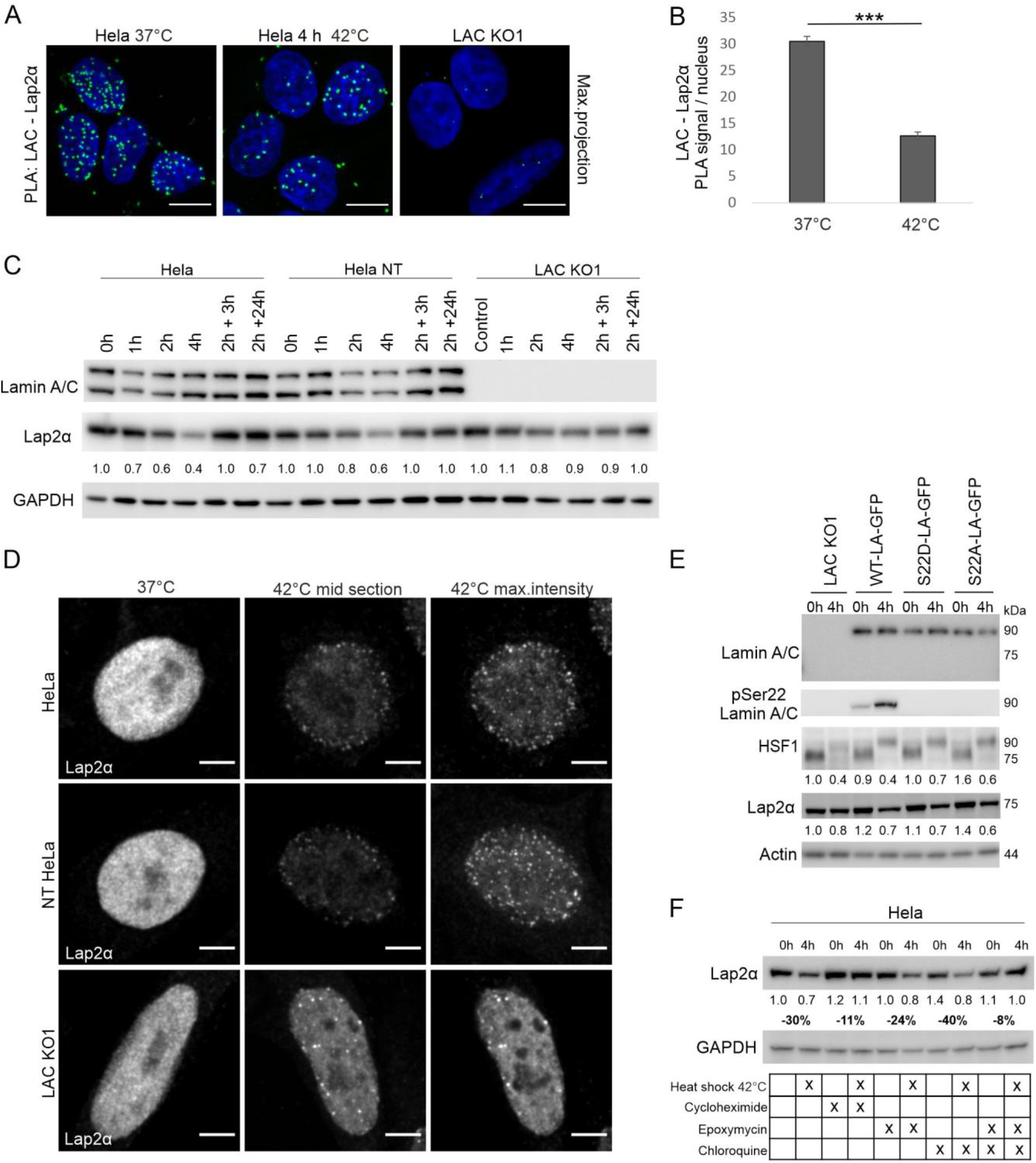
Lap2α is degraded in a lamin A/C dependent manner under HS. **A)** Proximity ligation assay (PLA) with lamin A/C and Lap2α antibodies was carried out on control and heat-shocked cells. Maximum projections of confocal images are shown. Scale bar 10 μm. Lamin A/C KO1 cells were used as a negative control. **B)** Quantification of PLA signals per nucleus (N=200, three individual experiments). **C)** Western blot analysis showing lamin A/C and Lap2α levels in HeLa, NT Hela, and LAC KO1 cells at different time point under HS and at the recovery. The average numerical values of signal intensities relative to the loading control (GAPDH) are shown below the blot. Each cell line has been normalized to its own control sample to highlight the change of Lap2α protein level between the timepoints (N=3). **D)** Confocal microscopy images showing Lap2α aggregation upon 4 h heat shock in Hela, NT Hela, and LAC KO1 cells. Scale bar 5 μm. **E)** Western blot analysis of LAC KO1 cells transfected with different GFP-tagged lamin A vectors and detected with pSer22 lamin A, HSF1, and Lap2α antibodies. The average numerical values of signal intensities relative to the loading control (actin) are shown below the blot (N=3). **F)** Western blot analysis of Lap2α protein level in heat-shocked parental Hela cells treated with either 10 μM CHX, 10 μM EPO, 10 μM CQ or both EPO and CQ. The average numerical values of signal intensities relative to the loading control (GAPDH) and percentage of intensity change between control and heat-shocked cells are shown below the blot (N=2). Data is expressed as mean ± s.e.m, ***p < 0.001.

Since HS-induced Lap2α downregulation could be due to reduced protein synthesis or degradation of the protein, we next exposed HeLa cells to HS in the presence of 10 μM cycloheximide (CHX, an inhibitor of protein synthesis), 10 μM epoximycin (EPO, a proteosomal inhibitor), and/or 10 μM chloroquine (CQ, an inhibitor of autophagy) (Fig. 6E). Surprisingly, Lap2α was not downregulated or aggregated in heat-shocked cells under CHX treatment. Since CHX inhibits all protein synthesis, it is possible that proteins required for Lap2α degradation were not produced. This also points out that Lap2α downregulation is due to degradation of the protein rather than inhibition of protein synthesis. Lap2α was equally downregulated upon HS in the presence of EPO or CQ. However, in the presence of both EPO and CQ, Lap2α protein level remained stable. These results suggest that under prolonged HS, Lap2α is degraded possibly through both proteasomes and autophagy, and the degradation is impaired in lamin A/C KO cells.

### Dermal patient fibroblasts with a LMNA mutation are more sensitive to HS

To study whether aforementioned changes in Lap2α take place also in normal diploid cells, control and patient fibroblasts with p.S143P *LMNA* mutation were tested. Similar to Hela cells, Lap2α was downregulated in heat-shocked control fibroblasts (Fig. 7A; p < 0.05). However, in the patient cells Lap2α downregulation was delayed and evident only after 4 h HS and at the recovery (Fig. 7A; p < 0.05). Further analysis revealed that Lap2α was downregulated in parallel with reduced proliferation marker Ki-67 expression (Fig. 7B,C, S7). In the control cells, the number of Ki-67 positive cells decreased from original 35% to 7% during 4-hour HS while in the patient cells a significantly smaller reduction from 41% to 28% was observed (Fig. 7C; p < 0.001 and p < 0.05, respectively). Correspondingly, there was significantly more mitotic cells among patient cells compared to controls during HS (Fig. 7D, p < 0.05). These results suggest that HS-induced cell cycle arrest to G0 is impaired in lamin A/C mutant cells. Additionally, there was significantly more cleaved poly (ADP-ribose) polymerase-1 (PARP-1) in the patient cells throughout the experiment, potentially indicating caspase activity and increased DNA damage in these cells (Fig. 7A). Cell viability was further analyzed with a cell count assay. HS at 42°C did not affect the viability of either cell line (Fig. S7). However, HS at 44°C reduced the survival of patient cells (presumably due to increased cell death) within the first 48 hours of recovery (Fig. 7E; p < 0.05). Both the cell lines eventually recovered from the HS and started to proliferate after 96 h. Altogether, these results suggest that patient cells have decreased capacity to undergo cell cycle arrest which may eventually lead to reduced cell survival at the recovery.

**Figure 7.**
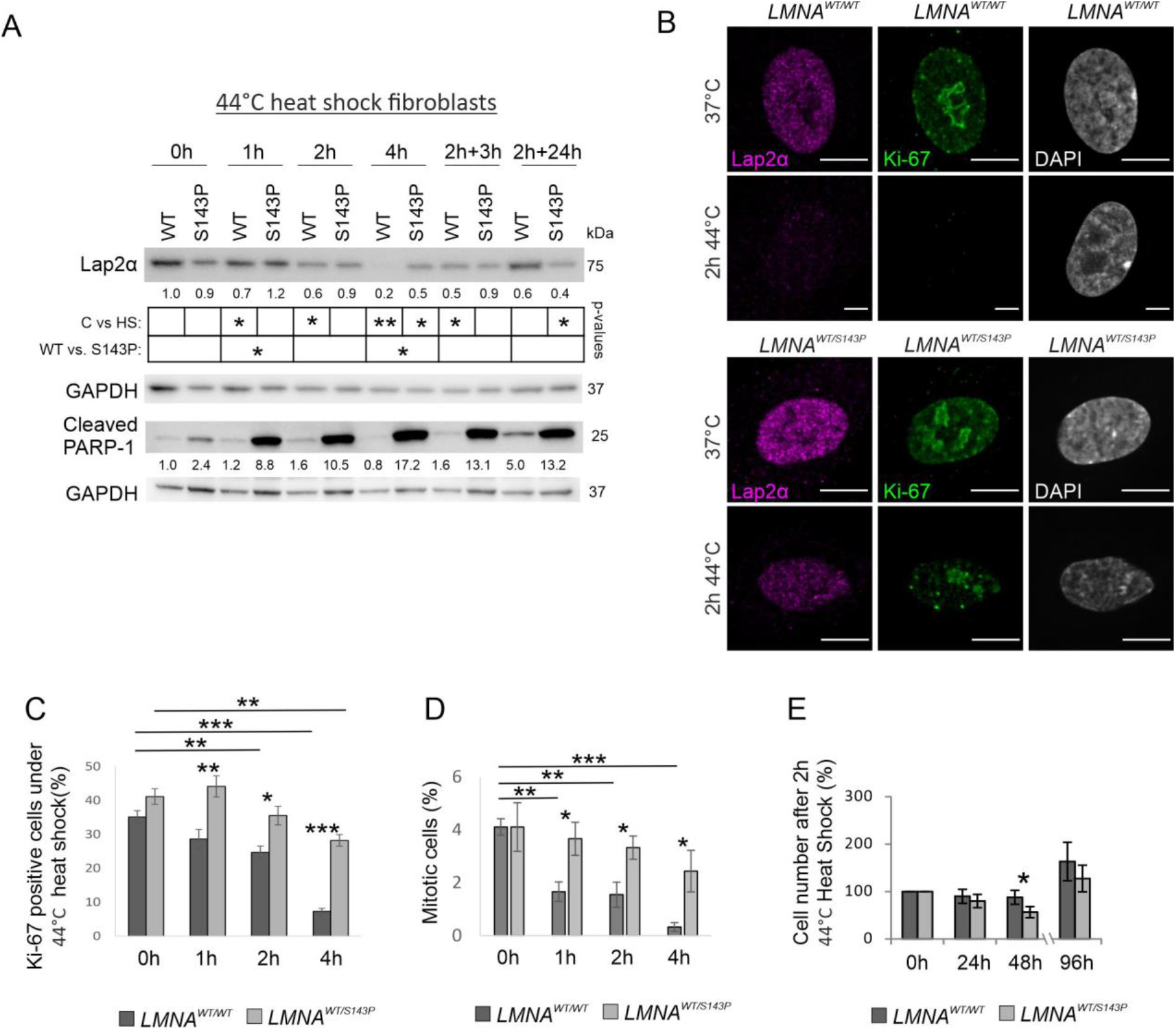
*LMNA* mutant cells are hypersensitive to severe HS. **A)** Western blot analysis of *LMNA^WT/WT^* and *LMNA^WT/S143P^* fibroblasts as detected with Lap2α and cleaved PARP-1 antibodies. The average numerical values of signal intensities relative to the loading control (GAPDH) are shown below each blot (N=3-5). **B)** Representative confocal images of *LMNA^WT/WT^* and *LMNA^WT/S143P^* cells cultured in normal conditions or exposed to 44°C for 2 hours prior to fixation. The cells were stained for Lap2α (magenta), Ki-67 (green) and DAPI (grey). **C)** The percentage of Ki-67 positive cells at 37°C and after 1- – 4-h at 44°C (N=1200). **D)** The percentage of mitotic cells at 37°C and after 1- to 4-h HS at 44°C (N=900). **E)** Cell numbers counted after 2-h HS at 44°C (time point 0 h) and after 24-, 48-, and 96-h recovery period at 37°C (N=6). Data is expressed as mean ± s.e.m, *** p < 0.001, ** p < 0.01, * p < 0.05.

## DISCUSSION

In the current study, we describe a previously unreported and evolutionary conserved post-translational modification of the lamin A/C amino-terminal head upon HS. Lamin A/C was gradually phosphorylated at Ser22 in both normal diploid and transformed human and mouse cell lines, as well as in *D. melanogaster* flies *in vivo.* Consequently, the phosphorylated pool of lamin A/C relocated into nucleoplasm and the adapted nuclei became more spherical especially under severe HS. On the other hand, lamin A/C KO and mutant cells were more vulnerable to HS indicating that normal filamentous lamina, formed by unphosporylated lamins, is equally important for nuclear shape maintenance and for regulation of cell cycle through Lap2α in heat-shocked cells. All these results suggest that the nuclear lamina is dynamically modified under HS to preserve cell homeostasis.

### The effects of HS-induced phosphorylation on lamin A/C

In early studies, Krachmarov & Traub (1993) reported heat-induced stabilization of nucleoskeleton and dephosphorylation of lamin A/C in Ehrlich Ascites tumor cells after HS. Additionally, Smith et al. (1987) reported that in *Drosophila* Schneider 2 cells lamin Dm2 is dephosphorylated to lamin Dm1 under HS. In contrast to these studies on the overall phosphorylation state, we focused on specific lamin A/C phospho-epitopes and our results are thus not necessarily in contradiction with the previous observations (although we found no clear evidence for general dephophorylation of lamin A/C upon HS in LC-MS/MS analysis).

Ser22 is a canonical mitotic phosphorylation site but accumulating data shows that Ser22 is also phosphorylated in interphase cells upon low mechanical stress and on soft matrices leading to increased elasticity of the nuclear lamina and improved cellular adaption (Bainer & Weaver, 2013; Buxboim et al., 2014). Our results are in line with the literature as we demonstrated that lamin A/C phosphorylation increases the nucleoplasmic localization of lamin A/C and simultaneous rounding off a nucleus in response to HS (Fig. 2C-I). Hence, the pSer22 response causes a partial change in phosphorylation stoichiometry, which is not sufficient to induce full disassembly but rather a partial shift from the assembled to the nucleoplasmic lamin pool, which is likely to serve the regulation of nuclear structure and properties to facilitate nuclear adaptation during the ongoing stressful conditions.

The nuclear rounding during HS is presumably a consequence of cell shrinking. Gungor et al. (2014) reported a drastic reduction of F-actin expression and cell rounding/shrinking in response to severe heat treatment (43°C). The resulting reduced cytoskeletal tension also decreases tension on the nucleus and facilitates its rounding through lamin A/C phosphorylation (Buxboim et al., 2014). In accordance with the assumption that nuclear rounding is due to phosphorylation-mediated effects, we found that nuclear rounding of fibroblast was prevented when lamin A/C phosphorylation was inhibited (Fig. 2, S3).

More recently, it was reported that in interphase cells pSer22 lamin A/C may act as a transcriptional activator by binding gene enhancer domains in the nuclear interior, and some of the binding sites were altered in dermal fibroblasts obtained from HGPS patients (Ikegami et al., 2020). Transcriptional response to HS results in the repression of thousands of genes (Mahat et al., 2016) and this is likely to explain the increased intensity of heterochromatin detected in healthy fibroblasts during HS (Fig. 3D). The enhanced heterochromatin intensity also correlated with pSer22 lamin A/C status (Fig. 3E). However, we did not observe a similar heterochromatin increase in the patient cells (Fig. 3E). Considering the findings reported by Ikegami et al. (2020) it is tempting to speculate that phosphorylated lamin A/C contributes to HS-induced gene expression, similar to HSF1, and further studies on these mechanisms are warranted.

We also noticed that lamin A/C was constantly more phosphorylated at Ser22 in the patient fibroblasts compared to control cells. Since p.S143P mutant lamin A/C is incapable of forming normal filaments and primarily nucleoplasmic (West et al., 2016), it may be more accessible by kinase for phosphorylation in these cells (Fig. 2A, B). As Ser22 phosphorylation is known to increase the mobility of lamin A (Kochin et al., 2014), the increased phosphorylation of lamin A/C may further reduce the stability of lamina in the patient cells.

### Multiple kinases may phosphorylate lamin A/C under HS

To determine the kinases responsible for the HS-induced phosphorylation of lamin A/C Ser22, we inhibited several different kinases that have previously been shown to phosphorylate lamin A/C (reviewed by Liu and Ikegami, 2020). Both MAPK and PKC inhibitors partially abolished lamin A/C phosphorylation, while the pan-inhibitor (STA) had the most significant effect suggesting that several kinases may be involved. Our results are consistent with previous screening studies showing that lamin A/C are ERK2 substrates (Carlson et al., 2011). There is also evidence that lamin A is phosphorylated at Ser268 by PKC to regulate nuclear size (Edens et al., 2017). Interestingly, we also detected the phosphorylation of Ser268 in LC-MS/MS analysis after 2- and 4-h HS but not in control cells (Table 1S). Therefore, it is possible that phosphorylation of lamin A/C at Ser268 contributes to modulation of nuclear size upon HS. Since at least 25 residues in lamin A/C are known to be phosphorylated during interphase and the degree of phosphorylation at each site may vary (Liu & Ikegami, 2020), one still needs to be cautious when estimating the biological effect of a single residue phosphorylation.

### The effects of Lap2α degradation on cell survival upon HS

Our results also revealed that Lap2α is degraded under HS and the degradation is impaired in lamin A/C KO cells (Fig. 6C, S5). Transfections with WT-LA, S22D-LA, and S22A-LA restored the degradation of Lap2α (Fig. 6E), which suggest that Lap2α is degraded in lamin A/C dependent manner, but is independent of lamin A/C Ser22 phosphorylation status. A similar downregulation was detected in primary human skin fibroblasts derived from a healthy control and a DCM patient, carrying the p.S143P *LMNA* mutation (Fig. 7A). We also noticed that Lap2α is degraded simultaneously with Ki-67 in control fibroblasts under severe HS but such degradation appeared delayed in the *LMNA* mutant patient fibroblasts (Fig. 7B-C). Previous data has shown that Lap2α is required for cell proliferation and is degraded upon cell cycle arrest in normal human fibroblasts (Pekovic et al., 2007). Based on these results, we conclude that lamin mutant fibroblasts have decreased capacity to undergo cell cycle arrest to G0 under HS. Similar results have been reported in *Lmna^−/−^* MEFs, that showed diminished cell cycle arrest in response to γ irradiation-induced DNA damage (Johnson et al., 2004). Continued cell cycle under stressful conditions typically leads to accumulating DNA damage and eventually cell death through apoptosis. Accordingly, PARP-1 was increasingly cleaved in lamin mutant patient cells under 44°C HS and the cells showed reduced survival at the recovery when compared to control cells (Fig. 7E). In support, hypersensitivity to HS has previously been reported in dermal fibroblasts obtained HGPS patients carrying the G608G mutation in *LMNA* gene. These cells showed dysmorphic nuclei and delayed recovery after 30 min HS at 45°C (Paradisi et al., 2005). Extensive nuclear deformations were also detected in heat-shocked dermal fibroblasts from FPLD patients carrying the R482Q mutation in *LMNA* gene (Vigouroux et al., 2001). Altogether these results indicate that in heat-shocked cells, where the lamina is rapidly remodeled through phosphorylation, the disease-linked mutations in lamin A/C cause nuclear instability and may sensitize cells to cell death.

In conclusion, we discovered a previously unknown and evolutionary conserved mechanism that regulates lamin A/C dynamics in heat-shocked cells. The degree of lamin A/C Ser22 phosphorylation appears tightly regulated and correlates with duration and severity of HS suggesting that the ratio of phosphorylated and unphosphorylated lamin A/C is critical for maintenance of proper lamina elasticity/stiffness, depending on prevailing circumstances. Our results also show that HS-induced modifications in the lamina are very similar to those induced by mechanical cues, such as matrix elasticity (Buxboim et al., 2014), and it seem plausible that Ser22 phosphorylation is a consequence of cell shrinking and nuclear rounding observed in heat-shocked cells. Which kinase is responsible for Ser22 phosphorylation and whether pSer22-lamin A/C also plays a role in HS-induced gene expression remains to be elucidated in follow-up studies.

## MATERIALS AND METHODS

### Cell lines, cell culture and transfections

HeLa cervical carcinoma cells were grown under a humidified 5% CO2 atmosphere at 37°C in Dulbecco’s modified Eagle’s medium (DMEM, Lonza, Alpharetta, GA, USA) supplemented with 10% fetal bovine serum (Thermo Fisher Scientific, Waltham, MA, USA) and penicillin/streptomycin/glutamine (Thermo Fisher Scientific). HSF1 silenced HeLa cells (ShHSF1) were kindly provided by Professor Lea Sistonen (Åbo Akademi University, Turku, Finland). Primary human and mouse fibroblasts were cultured in Minimum Essential Media (MEM, Thermo Fisher Scientific) supplemented with 15% FBS, antibiotics (penicillin, streptomycin), and 5% Non-Essential Amino Acids (Thermo Fisher Scientific). The use of patient fibroblasts was approved by the Ethics Committees of the Hospital District of Helsinki and Uusimaa (HUS 387/13/03/2009 and HUS/1187/2019). All procedures were undertaken with informed consent and according to the principles expressed in the Declaration of Helsinki.

HeLa cells and mouse fibroblasts were heat shocked under a humidified 5% CO2 atmosphere at 42°C for 1, 2, or 4 h and left to recover at 37°C for 3 or 24 h after 2-h HS. Human fibroblasts were heat shocked either at 42°C or 44°C. For cell cycle synchronization 10 μg/ml aphidicolin (Sigma-Aldrich, Saint Louis, MO, USA) was used 24 h prior to HS. For kinase inhibition 100 nM flavopiridol (Selleckhem, Houston, TX, USA), 1 μM roscovitine (Selleckhem), 10 μM U0126 (Selleckhem), 10 μM Go6976 (Selleckhem), 10 μM triciribine (API-2, Selleckhem), or 200nM staurosporine (STA, Selleckhem) was used 24 h prior to exposure to HS. For degradation analysis we exposed HeLa cells to HS in the presence of 10 μM cycloheximide (CHX, Sigma-Aldrich), 10 μM epoximycin (EPO, Sigma-Aldrich), and 10 μM chloroquine (CQ, Sigma-Aldrich). For control cells, equal concentrations of solvent (DMSO) was added into culture media.

Lamin A/C KO HeLa cells were transfected with Trans-IT HelaMONSTER (MirusBio, Madison, WI, USA) according to the manufacturer’s protocol and used for experiments 48 h after the transfection. Plasmids encoding either wild-type EGFP-tagged lamin A (pEGFP-C1-LA), phosphomimetic (pEGFP-C1-LA-S22D), or phospho-deficient (pEGFP-C1-LA-S22A) mutant forms of lamin A were cloned as described elsewhere (Kochin et al., 2014). Human fibroblast count was determined after 2-h HS (44°C) at 0, 24, 48, and 96-h time points. Attached cells were trypsinized at each time point, collected and counted with cell counting slides (BioRad, Hercules, CA, USA).

### CRISPR/Cas9 *LMNA* KO Hela

Lamin A/C knock-out (KO) HeLa cells were established with CRISPR/Cas9 technology at Turku Bioscience Genome Editing Core. A two-component CRISPR system was used to generate *LMNA* KO cells (Adli, 2018). *LMNA* sgRNAs (seq#1-CCAGAAGAACATCTACAGTG, seq#2-TGAAGAGGTGGTCAGCCGCG, seq#3-ATGCCAGGCAGTCTGCTGAG were selected using DeskGEN platform and cloned according to Feng Zhang lab protocol. Both *LMNA* sgRNA#1 and #2 generated full KO clones.

Separate lentivectors containing spCas9 (lentiCas9-Blast a gift from Feng Zhang (Addgene plasmid # 52962) and sgRNA (lentiGuide-Puro a gift from Feng Zhang (Addgene plasmid # 52963) were produced in 293FT packaging cell line by transient cotransfection. Shortly, 40-70% confluent HEK 293FT cells were used for transfections with 14 μg of transfer vector, 4 μg of packaging vector psPAX2 (gift from Didier Trono (Addgene plasmid # 12260)), 2 μg envelope vector pMD2.G (gift from Didier Trono (Addgene plasmid # 12259)) mixed in 0.45 mL water, 2.5M CaCl2, and 2x HeBS (274 mM NaCl, 10 mM KCl, 1.4 mM Na2HPO4, 15 mM D-glucose, 42 mM Hepes, pH 7.06) per 10 cm dish. Before adding to the cells, the DNA-HeBS mix was incubated for 30 min at RT. After overnight incubation medium with DNA precipitate was gently removed from the cells and replaced with a full fresh medium. Media containing viral particles was collected after 72 h, spun at 300 rpm for 5 min at RT to remove cell debris, filtered through 0.45 μm PES filter, and concentrated by ultracentrifugation for 2 h at 25,000 rpm, 4°C (Beckman Coulter). The pellet containing lentiviral particles was suspended in the residual medium, incubated for ~2 h in +4°C with occasional mild vortex, aliquoted, snap-frozen and stored in −70°C. P24 ELISA measured physical lentiviral titer with a serial dilution of virus stock according to manufacturer protocol.

To generate *LMNA^−/−^* clones in HeLa cells, 1e+05 cells were seeded on a 24-well plate. The next day the cells were transduced with Lenti-Cas9 (MOI 1, 3, 6), and 72 hours later, 8 μg/mL of Blasticidin was applied to select Cas9 expressing cells. Cells transduced with the smallest amount of Lenti-Cas9 particles that survived after the control well were proceeded to the next step. In the second stage, the mixed pool of stably expressing Cas9 cells was transduced with Lenti sgRNA vectors (MOI 6,9, 12), and 72 hours later, 1 μg/ul Puromycin was applied on cells to select double-positive Cas9+/Lenti sgRNA+ cells. Based on Western blot results, cell populations showing the highest reduction in lamin A/C protein levels were single sorted (Sony SH800 cell sorter, Sony Biotechnology Inc) and re-grown into a clonal cell population. On average, about 20 clones per sgRNA population were screened with Western blotting and Sanger sequencing to confirm full knockout status.

### *Drosophila melanogaster* husbandry, treatments and analysis

Canton-S wild type flies were a gift from Dr. Pascal Meier, Institute of Cancer Research, London UK. Flies were maintained in vials at 22°C on a 12-hour light/dark cycle on Nutri-Fly cornmeal medium (Nutri-fly BF, Dutscher Scientific, France). Adult flies were used in the experiments. HS was induced in the flies by incubating the vials (20-30 flies per vial) at 37°C for 30 min, followed by 3- or 7-h recovery periods at 22°C. Ten flies per sample were kept at −20°C for 5 min to euthanize the flies. Using an electronic pestle, the flies were homogenized in lysis buffer (50 mM Tris [pH 7.5], 150 mM NaCl, 1% Triton X-100, 1 mM EDTA, 10% glycerol) with 1x protease and phosphatase inhibitors (Thermo Scientific). The lysates were kept on ice for 10 min. To remove exoskeleton and fat from the lysates, the samples were centrifuged for 10 min at 12000 rpm at 4°C. The clear supernatant was transferred into new microtubes, avoiding the debris at the bottom and the fat layer on the top. The lysate was cleared by centrifuging again for 10 min at 12000 rpm at 4°C. The clear supernatant was transferred into new microtubes and mixed with 4x Laemmli-β-mercaptoethanol to a concentration of 1x. Samples were incubated at 95°C for 10 min and separated on 10% SDS-PAGE gel and transferred to a nitrocellulose membrane. *Drosophila* lamin C and phosphorylated lamin C were detected with mouse anti-lamin C antibody (1:1000, LC28.26, DSHB, Iowa City, IA, USA) and rabbit monoclonal phospho-lamin A/C (Ser22) (1:1000, D2B2E, Cell Signaling Technology, Danvers, MA, USA), respectively. Actin was used as a loading control and was detected with goat polyclonal actin antibody (1:2000, C11, sc-1615, Santa Cruz Biotechnology, Dallas, TX, USA). The secondary antibodies were goat anti-mouse IgG-HRP (sc-2005), goat anti-rabbit IgG-HRP (sc-2004) and donkey anti-goat IgG-HRP (sc-2020) (all from Santa Cruz Biotechnology) at 1:5000. The ECL advanced Western blotting detection kit (RPN 3243, Sigma Aldrich) was used for signal detection.

### Immunoblotting

Unheated and heated samples were lysed in mammalian protein extraction reagent (M-PER, Thermo Fisher Scientific) complemented with 1x protease and 1x phosphatase inhibitors (Thermo Fisher Scientific). Cell lysates were mixed with 4x Laemmli sample buffer and run on a 4–20% gradient gel (BioRad) and transferred to nitrocellulose filter (BioRad). The primary antibodies used were mouse monoclonal lamin A/C (1:10,000, 5G4, kindly provided by Prof. Robert D. Goldman, Northwestern University, USA), rabbit polyclonal heat shock factor 1 (1:1000, ADI-SPA-901, Enzo Life Science, Farmingdale, NY, USA), rat monoclonal heat shock factor 1 (1:1000, 10H8, Stress Marq Bioscience inc., Victoria, Canada), rabbit monoclonal phospho-lamin A/C serine 22 (1:1000, D2B2E, Cell Signaling Technology), rabbit polyclonal phospho-lamin A/C serine 392 (1:1000, ab58528, Abcam), rabbit polyclonal AKT (pan) (1:1000, C67E7, Cell Signaling Technologies), rabbit monoclonal pAKT T303 (1:1000, D6F8, Cell Signaling Technologies), rabbit polyclonal ERK1 (1:500; K-23, Santa Cruz Biotechnology), rabbit polyclonal ERK2 (1:500, c-14, Santa Cruz Biotechnology), rabbit monoclonal pERK1/2 (1:1000, D13.14.4E, Cell Signaling Technologies), rabbit monoclonal cleaved PARP-1 (1:1000, E-51, Abcam, Cambridge, UK), rabbit monoclonal Caspase-3 (1:1000, 8G10, Cell Signaling), rabbit monoclonal cleaved caspase-3 (1:1000, Asp175, Cell Signaling), rabbit monoclonal Lap2α (1:5000, 245/2, kindly provided by Prof. Roland Foisner, University of Vienna), HSP70/HSP72 mouse monoclonal (1:1000, C92F3A-5, Enzo life sciences), HSC70/HSP73 rat monoclonal (1:1000, 1B5, Enzo life sciences), HRP-conjugated anti-GAPDH (1:5000, ab9385, Abcam) and mouse monoclonal anti-actin (1:1000, AC-40, Sigma-Aldrich). Secondary antibodies were HRP-conjugated donkey anti-rabbit-IgG, sheep anti-mouse-IgG and anti-rat-IgG (all from GE Healthcare, Chicago, IL, USA). The antibodies were detected with Enhanced Chemiluminescence kit (Thermo Fischer Scientific).

### Immunofluorescence and Microscopy

Cells grown on coverslips were fixed in 10% formalin for 10 min, permeabilized with 0.1% Triton X-100 for 10 min and blocked with 1% BSA in TBST for 30 min. The primary antibodies used were mouse monoclonal lamin A/C (1:10000, 5G4, kindly provided by prof. Robert D. Goldman, Northwestern University, USA), mouse monoclonal lamin A/C (1:100, 4C11, Cell Signaling Technologies), rabbit monoclonal phospho-lamin A/C serine 22 (1:400, D2B2E, Cell Signaling), rabbit polyclonal HSF1 (1:400, ADI-SPA-901, Enzo), rat monoclonal HSF1 (1:400, 10H8, StressMarq Bioscience Inc.), rabbit monoclonal Lap2α (1:1000, 245/2, kindly provided by Prof. Roland Foisner, University of Vienna), and mouse monoclonal Ki-67 (1:200, MIB-1, DAKO, Denmark). The secondary antibodies were donkey anti-rabbit IgG conjugated to Alexa Fluor 488, donkey anti-mouse IgG conjugated to Alexa Fluor 555, and chicken anti-rat IgG conjugated to Alexa Fluor 647 (all Molecular Probes, 1:200, Eugene, OR, USA). ProLong Diamond Antifade Mountant with DAPI was used to visualize DNA (Thermo Fisher Scientific). Dualink proximity ligation assay (PLA) was conducted according to the manufacturer’s protocol (Sigma-Aldrich).

The spinning disk confocal microscope used was a 3i Marianas with Yokogawa CSU-W1 scanning unit on an inverted Zeiss AxioObserver Z1 microscope, controlled by SlideBook 6 software (Intelligent Imaging Innovations GmbH, Göttingen, Germany). The objective used was 63x/1.4 oil. Images were acquired with ORCA Flash4 sCMOS camera (Hamamatsu Photonics, Hamamatsu, Japan). All the images were analyzed with ImageJ Fiji software (Schindelin et al., 2012). Confocal maximum projection images were used to analyze the quantity of PLA signals. Lap2α aggregate quantity and localization were analyzed from 3D confocal stacks using imageJ plugin NucleusJ (Poulet et al., 2015). Fluorescence intensities of lamin A/C within the lamina (L) and nucleoplasmic (N) regions were quantified and the ratio of fluorescence between the lamina and nucleoplasma were calculated as follows:

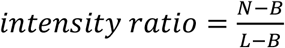

where B is background.

The shHSF1 samples were imaged using a Zeiss LSM880 confocal microscope with Airyscan mode. Images were taken using a 63x Zeiss C plan-Apochromat oil immersion objective (NA = 1.4) with an additional 1.8x zoom. Following imaging, the raw data was processed using the Airyscan processing module in the Zen blue software (Jena, Germany). Minor image quality adjustments were done using the ImageJ FIJI software.

### Phosphorylation analysis by mass spectrometry

Lamin A/C was immunoprecipitated from G1/S phase synchronized control and heat shocked (2 and 4 h at 42°C) HeLa cells and run on gel. Mass spectrometry analysis was performed at the Turku Proteomics Facility, University of Turku and Åbo Akademi University with the following protocol: Lamin A/C immunoprecipitate was in-gel digested using trypsin. Phosphopeptides were enriched by Thermo’s High-Select TiO2 Phosphopeptide Enrichment Kit according manufacturer’s protocol. The LC-ESI-MS/MS analyses were performed on a nanoflow HPLC system (Easy-nLC1200, Thermo Fisher Scientific) coupled to the Q Exactive HF mass spectrometer (Thermo Fisher Scientific, Bremen, Germany) equipped with a nano-electrospray ionization source. MS data was acquired automatically by using Thermo Xcalibur 3.1 software (Thermo Fisher Scientific). Data files were searched for protein identification using Proteome Discoverer 2.3 software (Thermo Fisher Scientific) connected to an in-house server running the Mascot 2.6.1 software (Matrix Science). Data was searched against a SwissProt database containing human protein sequences.

### Statistical analysis

Each experiment was repeated at least two times from samples that were prepared independently of each other. The results are presented as average values ± SEM. The statistical analyses were performed using Graphpad Prism 8, Microsoft Excel or R software. Datasets that followed normal distribution were analyzed with Student’s t-test or grouped two-way ANOVA followed by Tukey post-test. Pearson correlation coefficient was used to quantifie the relationship between two variables. Non-parametric datasets were analyzed with Mann-Whitney test. P-values ≤0.05 were considered statistically significant and are marked with symbols in the figures. The exact sample sizes and number of measurements for each experiment is indicated in the figure texts.

## Acknowledgments

Jenny Joutsen is thanked for her advice and support on HS experiment, Robert D. Goldman for kindly providing the lamin antibodies, Lea Sistonen for shHSF1 cell line, and Roland Foisner for lap2α antibody. Cell Imaging Core (Turku Centre for Biotechnology) is acknowledged for help with confocal microscopy and AFM, Turku Proteomics Facility for the mass spectrometry, and Turku Bioscience Genome Editing Core for CRISPER/Cas9 lamin A/C KO cell lines. We also thank the patients for donating their biopsies for research.

## Author contributions

L.V., E.H., J.E., A.M. and P.T. designed and performed research, and wrote the manuscript. J.G., M.H., J.I., G.W, A.P. and F.L. performed experiments. T.H. recruited patients and provided material. All co-authors approved the final version of the manuscript.

## References

Adli, M. (2018). The CRISPR tool kit for genome editing and beyond. Nature Communications, 9, 1911.

Åkerfelt, M., Morimoto, R. I., & Sistonen, L. (2010). Heat shock factors: Integrators of cell stress, development and lifespan. Nature Reviews Molecular Cell Biology, 11, 545–555.

Bainer, R., & Weaver, V. (2013). Strength under tension. Science, 341, 965–966.

Buxboim, A., Swift, J., Irianto, J., Spinler, K. R., Dingal, P. C. D. P., Athirasala, A., Kao, Y. R. C., Cho, S., Harada, T., Shin, J. W., & Discher, D. E. (2014). Matrix elasticity regulates lamin-A,C phosphorylation and turnover with feedback to actomyosin. Current Biology, 24, 1909–1917.

Carlson, S. M., Chouinard, C. R., Labadorf, A., Lam, C. J., Schmelzle, K., Fraenkel, E., & White, F. M. (2011). Large-scale discovery of ERK2 substrates identifies ERK-mediated transcriptional regulation by ETV3. Science Signaling, 4, rs11.

Dynlacht, J. R., Story, M. D., Zhu, W. G., & Danner, J. (1999). Lamin B is a prompt heat shock protein. In Journal of Cellular Physiology, 178, 28–34.

Edens, L. J., Dilsaver, M. R., & Levy, D. L. (2017). PKC-mediated phosphorylation of nuclear lamins at a single serine residue regulates interphase nuclear size in Xenopus and mammalian cells. Molecular Biology of the Cell, 28, 1389–1399.

Gungor, B., Gombos, I., Crul, T., Ayaydin, F., Szabó, L., Török, Z., Mátés, L., Vígh, L., & Horváth, I. (2014). Rac1 participates in thermally induced alterations of the cytoskeleton, cell morphology and lipid rafts, and regulates the expression of heat shock proteins in B16F10 melanoma cells. PLoS ONE, 9, e89136.

Haddad, N., & Paulin-Levasseur, M. (2008). Effects of heat shock on the distribution and expression levels of nuclear proteins in HeLa S3 cells. Journal of Cellular Biochemistry, 105, 1485–1500.

Ikegami, K., Secchia, S., Almakki, O., Lieb, J. D., & Moskowitz, I. P. (2020). Phosphorylated Lamin A/C in the Nuclear Interior Binds Active Enhancers Associated with Abnormal Transcription in Progeria. Developmental Cell, 52, 699–713.e11.

Johnson, B. R., Nitta, R. T., Frock, R. L., Mounkes, L., Barbie, D. A., Stewart, C. L., Harlow, E., & Kennedy, B. K. (2004). A-type lamins regulate retinoblastoma protein function by promoting subnuclear localization and preventing proteasomal degradation. Proceedings of the National Academy of Sciences of the United States of America, 101, 9677–9682.

Kline, M. P., & Morimoto, R. I. (1997). Repression of the heat shock factor 1 transcriptional activation domain is modulated by constitutive phosphorylation. Molecular and Cellular Biology, 17, 2107–2115.

Kochin, V., Shimi, T., Torvaldson, E., Adam, S. A., Goldman, A., Pack, C. G., Melo-Cardenas, J., Imanishi, S. Y., Goldman, R. D., & Eriksson, J. E. (2014). Interphase phosphorylation of lamin A. Journal of Cell Science, 127, 2683–2696.

Krachmarov, C. P., & Traub, P. (1993). Heat-induced morphological and biochemical changes in the nuclear lamina from Ehrlich ascites tumor cells in vivo. Journal of Cellular Biochemistry, 52, 308–319.

Lindquist, S. (1986). The Heat-Shock response. Annual Review of Biochemistry, 55, 115–91.

Liu, S. Y., & Ikegami, K. (2020). Nuclear lamin phosphorylation: an emerging role in gene regulation and pathogenesis of laminopathies. Nucleus, 11, 299–314.

Mahat, D. B., Salamanca, H. H., Duarte, F. M., Danko, C. G., & Lis, J. T. (2016). Mammalian Heat Shock Response and Mechanisms Underlying Its Genome-wide Transcriptional Regulation. Molecular Cell, 62, 63–78.

Paradisi, M., McClintock, D., Boguslavsky, R. L., Pedicelli, C., Worman, H. J., & Djabali, K. (2005). Dermal fibroblasts in Hutchinson-Gilford progeria syndrome with the lamin A G608G mutation have dysmorphic nuclei and are hypersensitive to heat stress. BMC Cell Biology, 6, 1–11.

Pekovic, V., Harborth, J., Broers, J. L. V., Ramaekers, F. C. S., Van Engelen, B., Lammens, M., Von Zglinicki, T., Foisner, R., Hutchison, C., & Markiewicz, E. (2007). Nucleoplasmic LAP2α-lamin A complexes are required to maintain a proliferative state in human fibroblasts. Journal of Cell Biology, 176, 163–172.

Poulet, A., Arganda-Carreras, I., Legland, D., Probst, A. V., Andrey, P., & Tatout, C. (2015). NucleusJ: An ImageJ plugin for quantifying 3D images of interphase nuclei. Bioinformatics, 31, 1144–1146.

Pradhan, R., Nallappa, M. J., & Sengupta, K. (2020). Lamin A/C modulates spatial organization and function of the Hsp70 gene locus via nuclear myosin I. Journal of Cell Science 133, jcs236265.

Schindelin, J., Arganda-Carreras, I., Frise, E., Kaynig, V., Longair, M., Pietzsch, T., Preibisch, S., Rueden, C., Saalfeld, S., Schmid, B., Tinevez, J. Y., White, D. J., Hartenstein, V., Eliceiri, K., Tomancak, P., & Cardona, A. (2012). Fiji: An open-source platform for biological-image analysis. Nature Methods, 9, 676–682.

Smith, D. E., Gruenbaum, Y., Berrios, M., & Fisher, P. A. (1987). Biosynthesis and interconversion of Drosophila nuclear lamin isoforms during normal growth and in response to heat shock. Journal of Cell Biology, 105, 771–790.

Swift, J., Ivanovska, I. L., Buxboim, A., Harada, T., Dingal, P. C. D. P., Pinter, J., Pajerowski, D., Spinler, K. R., Shin, J. W., Tewari, M., Rehfeldt, F., Speicher, D. W., & Discher, D. E. (2013). Nuclear Lamin-A Scales with Tissue Stiffness and Enhances Matrix-Directed Differentiation. Science, 341, 965–966.

Vigouroux, C., Auclair, M., Dubosclard, E., Pouchelet, M., Capeau, J., Courvalin, J. C., & Buendia, B. (2001). Nuclear envelope disorganization in fibroblasts from lipodystrophic patients with heterozygous R482Q/W mutations in the lamin A/C gene. Journal of Cell Science, 114, 4459–4468.

West, G., Gullmets, J., Virtanen, L., Li, S. P., Keinänen, A., Shimi, T., Mauermann, M., Heliö, T., Kaartinen, M., Ollila, L., Kuusisto, J., Eriksson, J. E., Goldman, R. D., Herrmann, H., & Taimen, P. (2016). Deleterious assembly of the lamin A/C mutant p.S143P causes ER stress in familial dilated cardiomyopathy. Journal of Cell Science, 129, 2732–2743.

Wu, C. (1995). Heat shock transcription factors: Structure and regulation. Annual Review of Cell and Developmental Biology, 11, 441–469.

Zhu, W. G., Roberts, Z. V., & Dynlacht, J. R. (1999). Heat-induced modulation of lamin B content in two different cell lines. Journal of Cellular Biochemistry, 75, 620–628.

